# Comparing evolvability and pleiotropy across environments

**DOI:** 10.1101/2022.04.01.485835

**Authors:** Jason Tarkington, Rebecca Zufall

**Affiliations:** University of Houston; Stanford University

## Abstract

The environment can play an important role in determining evolutionary outcomes (Reboud and Bell 1997; Stanton et al. 2000; Gresham et al. 2008; Cooper and Lenski 2010; Becks and Agrawal 2012; Bailey et al. 2015). Populations may increase in fitness more after evolution in one environment than in another (Hegreness et al. 2008). This may be because the distance between the ancestral genotype and the top of the locally accessible fitness peak may be greater in one environment or the other. Additionally, the rate at which mutations occur and populations move up the peaks could differ. For this reason, comparing evolvability across environments presents an interesting problem. Here, we show an environment impacts evolvability in two ways, directly by impacting the rate of adaptative evolution, and indirectly by limiting the range of fitness outcomes that are possible. We show that correlated responses are often highly idiosyncratic, due to variation in the range of possible fitness outcomes in the environment and differences in pleiotropic effects across environments but can also be predicted from the ancestral growth rate regardless of the environment in which populations evolve. Interestingly, we also show a negative correlation between increase in an environment X following evolution in environment Y and the increase in environment Y following evolution in environment X. These results highlight the necessity to measure fitness in both environments when comparing the evolvability or repeatability of evolution across environments.

## Introduction

We found that the relative increase in fitness of replicate populations evolved at 24°C and 37°C was dependent on the assay environment and the evolution environment. Here, we aim to test the generality of these findings and specifically whether assay environment could systematically bias estimates of the effect of evolution environment on fitness increases when correlated responses are not measured. We also assessed if ancestral fitness was predictive of future fitness increases whether evolution occurred in that environment or not (Figure 3), whether fitness increases in the evolution environment can be used to predict fitness increases in other environments (Figure 4), and whether correlated responses are symmetric with respect to the assay and evolution temperature (Figure 5). To address these issues, we evolved 14 initially identical populations of the ciliate *Tetrahymena thermophila* in a range of environments and measured the fitness increase of each population in each of the environments. We found that assay environment as well as evolution environment had a significant effect on the increase in fitness demonstrating that evolvability is often constrained by natural limitations on fitness in an environment and that this aspect of evolvability can operate independently of the evolution promoting propensity of the environment.

## Methods

### High and Low Temperature Adaptation

We allowed 12 populations from 3 starting genotypes (4 of each) to evolve for 4000 generations at 24°C and another set of 12 from the same 3 starting genotypes to evolve at 37°C (Tarkington and Zufall 2021). Growth rate was measured at both temperatures as evolution progressed (see Tarkington and Zufall 2021 for further details on growth rate measurement). For each population, growth rate measurements at either temperature taken between 3900-4100 generations were binned and the mean was calculated. An ANOVA was performed on this data testing the effect of assay temperature, evolution temperature, and genotype on growth rate increases.

### Novel Environment Adaptation

13 populations were founded from a clonal population of the laboratory strain SB210E. Each population was allowed to evolve in a unique nutrient rich culture media containing inhibitory levels of various organic or inorganic compounds. These included 5% glycerol, 3% ethanol, 3% ethanol no glucose, 1.5% bleach, 0.18% citric acid, 0.18% citric acid no glucose, 0.04M NaOH, 0.05M NaOH, 15 g/L acetate, 15 g/L CaNO3, and 25 g/L NaCl. In some cases, the original concentration was lower at the start of the evolution experiment and gradually increased as the population began to grow faster and survive in concentrations that were previously lethal. The starting concentration for each environment was 2% glycerol, 2% ethanol, 2% ethanol no glucose, 1.5% bleach, 0.12% citric acid, 0.12% citric acid no glucose, 0.02M NaOH, 15 g/L acetate, 15 g/L CaNO3, and 15 g/L NaCl. Populations were maintained by serial dilution in 10mL of the specific media for each population. Dilution factors were adjusted so that daily bottlenecks never fell below ∼20,000 cells and effective population size was maintained at ∼100,000 cells. Slower growing cultures experienced a larger bottleneck but didn’t grow as dense as faster growing cultures. All populations were maintained for at least 2000 generations in one of 11 unique environments. After at least 2000 generations of evolution, growth rate and maximum OD of each of the evolved populations and the ancestor (thawed from stock) were assayed in each of the 11 unique environments. At least 4 growth curves per population were obtained for each of the 11 assay environments. The mean growth rate and maximum OD were then calculated for each population and assay condition. For some unique environments the concentration of the inhibitory compound was reduced to allow for growth of the evolved populations. However, for some combinations of evolution and assay environment we were still unable to obtain good growth curves. Concentrations used in growth assays were 5% glycerol, 1.5% ethanol, 1.5% ethanol no glucose, 0.6% and 1.5% bleach, 0.162% citric acid, 0.162% citric acid no glucose, 0.025M NaOH, and 15 g/L Acetate.

## Results

### High and Low Temperature Adaptation

The relative and absolute increase in growth rate after 4000 generations of evolution was calculated with 95% confidence intervals for each combination of genotype, evolved environment, and assay environment. Populations evolved and assayed at 24°C had a greater relative increase in fitness than populations evolved and assayed at 37°C (187% vs. 120%), while at the same time populations evolved at 37°C increased more in relative fitness when assayed at either temperature (24°C assay: 191% vs. 187%, 37°C assay: 120% vs. 108%; figure 1). However, as we state above, directly comparing fitness trajectories of populations evolved and assayed in one environment to those evolved and assayed in another environment confounds assay temperature and evolution temperature. For example, this difference between populations evolved at either temperature is driven almost entirely by differences in assay temperature as we show below.

**Figure 1.**
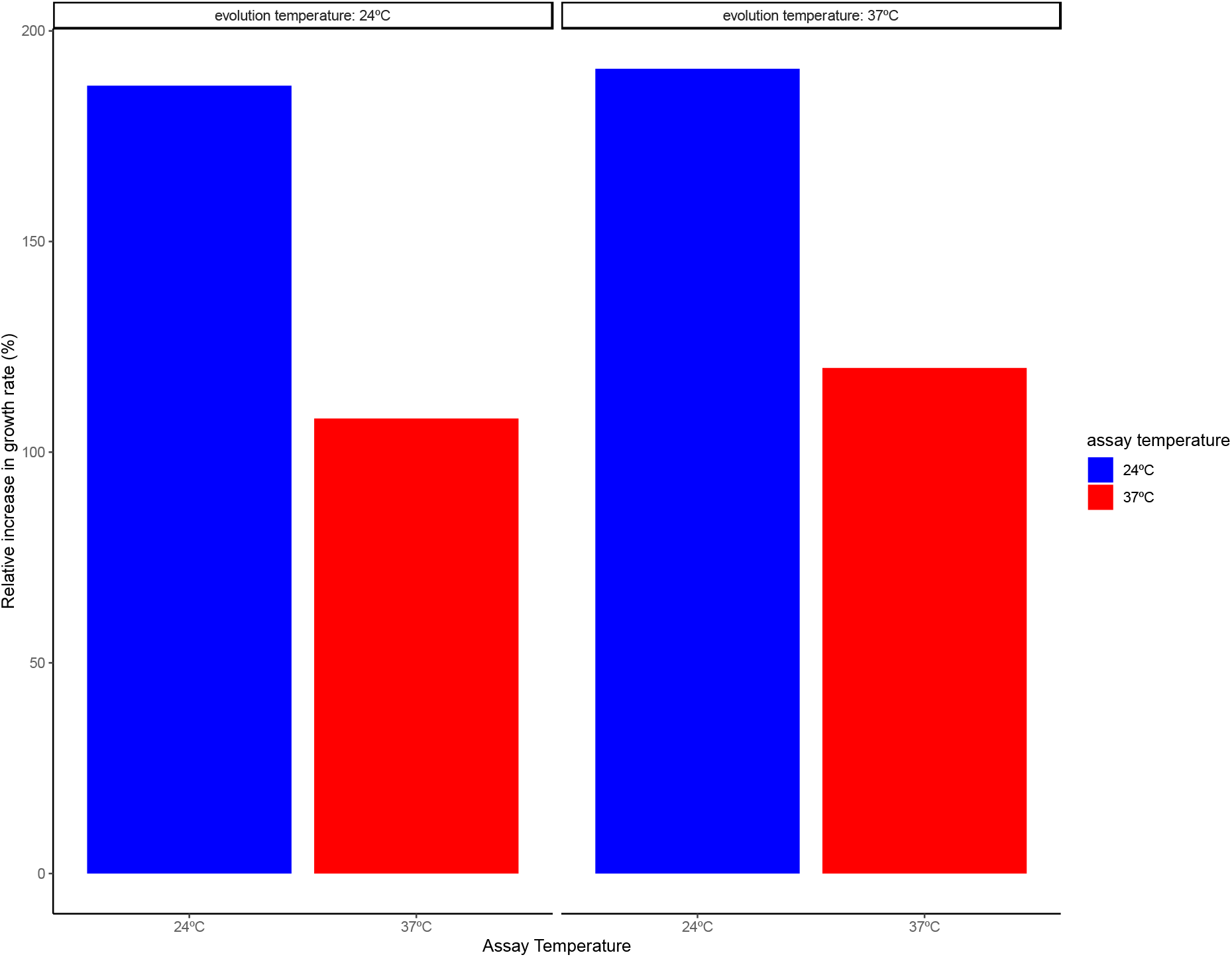
Relative increase in growth rate at 24°C (blue) and 37°C (red) of populations after evolution at either 24°C (left) or 37°C (right). Populations that evolved at 24°C increased more in relative growth at 24°C than those evolved at 37°C increased in relative growth rate at 37°C. However, the populations that evolved at 37°C have larger increases in relative growth rate at both temperatures (24°C assay: 191% vs. 187%, 37°C assay: 120% vs. 108%).

We performed two analyses to test the effects of genotype, evolution temperature, assay temperature, and 2-way interactions, on the absolute increase in growth rate and on the percentage increase. Each analysis included 48 data points corresponding to either the absolute increase or percentage increase in growth rate of each replicate population at either assay temperature. Results showed that assay temperature, but not evolution temperature, significantly affected both the percent increase (ANOVA: *F* (1,38) = 256.85, *P* < 0.0001) and the absolute increase (ANOVA: *F* (1,38) = 16.69, *P* = 0.0002) in growth rate. However, the absolute increase in growth rate was greater when assayed at 37°C regardless of the evolution temperature while the percent increase was greater when assayed at 24°C regardless of the evolution temperature. Unlike assay temperature the effect of evolution temperature was only marginally significant, however the effect was in the same direction, with the 37°C populations increasing more in absolute (ANOVA: *F* (1,38) = 2.262, *P* = 0.1408) and relative fitness (ANOVA: *F* (1,38) = 2.96, *P* = 0.0934).

After 4000 generations the 37°C evolved populations also had significantly higher growth rates than population evolved at 24°C (ANOVA: *F* (1,38) = 8.42, *P* = 0.0061) indicating more evolution at the hotter temperature despite a greater relative increase in fitness in the evolution environment after evolution at the colder temperature. While these results differ at other timepoints, we choose to report the results after 4000 generations to highlight the importance of measuring correlated responses when comparing evolvability across environments.

### Novel Environment Adaptation

The mean relative increase in growth rate (((evolved growth rate - ancestral growth rate) / ancestral) x 100) of each population assayed in each environment is shown in figure 2. Some of the evolved populations increase more in all environments (glycerol) while others see little increase in any environments (EtOH no gluc.). At the same time some of the environments (Acetate, NaOH) have large increases in fitness regardless of evolution environments while in others (etoh no gluc., glycerol) fitness seems to be constrained and experiences limited increases for all of the evolved populations. In fact, as we would expect based on diminishing returns epistasis, ancestral fitness was a good predictor of the total increase in fitness in that environment regardless of the evolution environment (figure 3). ANOVA results confirmed that both evolution environment (ANOVA: F(12,133) = 4.767, P < 0.0001) and assay environment (ANOVA: F(9,136) = 3.063, P < 0.0006) have a significant effect on the relative increase in growth rate. Similar results were obtained for the mean relative increase in maximum OD (ANOVA: evolution environment - F(12,132) = 4.734, P < 0.0001 and assay environment - F(9,135) = 5.960, P < 0.0001). We expected the largest increase in fitness to occur in environments in which evolution occurred, however we found that the three largest fitness increases we measured did not happen in the environment in which that population evolved.

**Figure 2.**
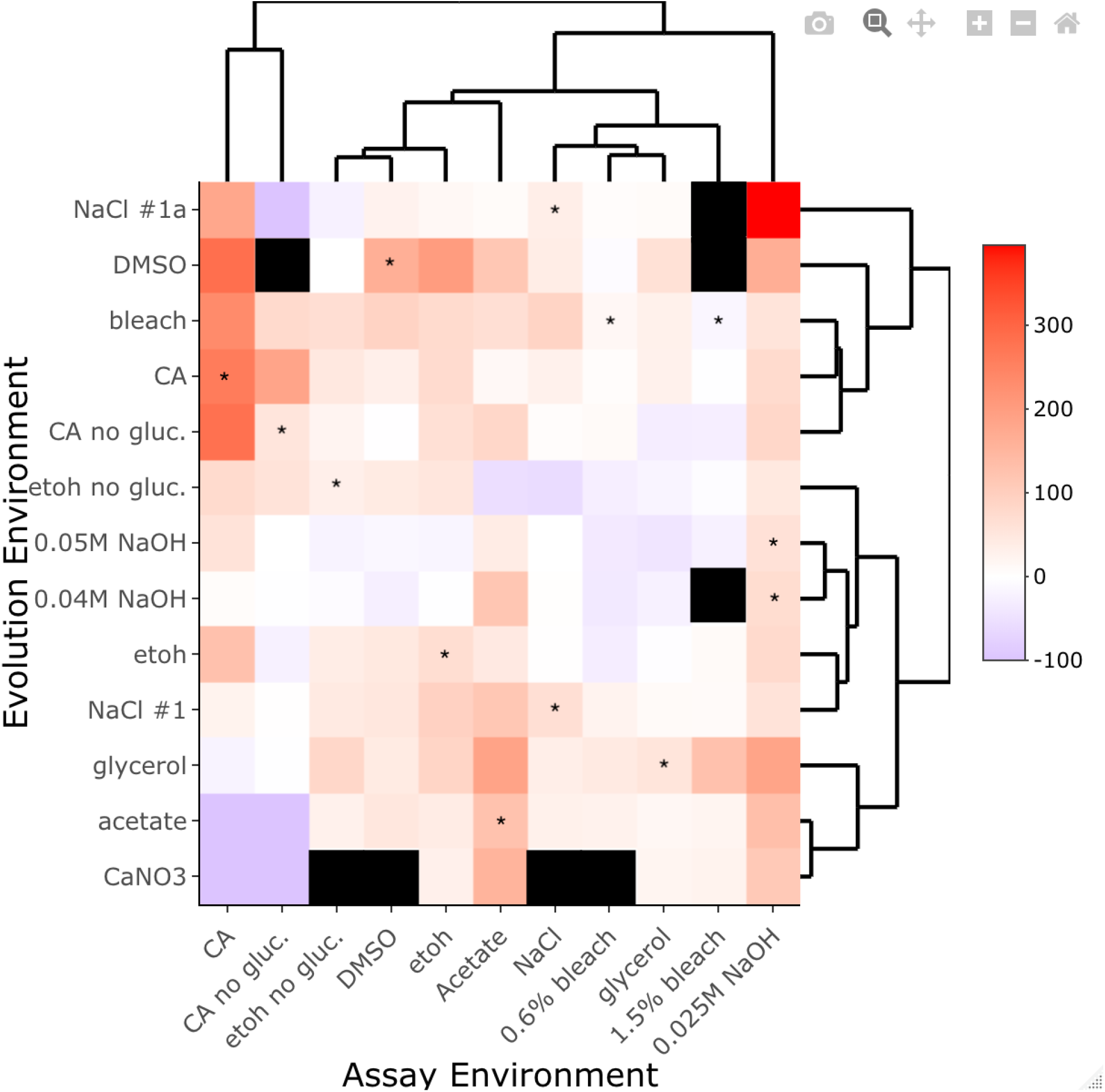
Heat map showing the relative (%) increase in growth rate in 11 environmental assay conditions of the 13 populations evolved under various environmental conditions. Each square shows the mean fitness increase in an assay environment for a population evolved in 1 of the 11 different environments. Black squares are missing data. Evolution and assays environments were clustered using the ‘heatmaply’ package in R.

We see several cases in which evolved populations have lower fitness in an environment than their ancestor (Fig. 2), however because we lack replicates within an evolution environment we cannot say anything about the systematic evolution of trade-offs between any two environments. Instead increases in growth rate in the evolved environment can be plotted against increases in growth rate in other environments to get a sense of the pattern of correlated responses that might emerge across a variety of environmental conditions. Figures 4 shows the mean relative increase in growth rate and maximum OD plotted against the increase in the other environments. We expected most populations to increase in fitness more in the environment in which they evolved than in other environments, however for r-max we found 43/124 of the points fall above the x=y line indicating many instances of a larger increase in growth rate in another environment. For maximum OD we observed 35/115 instances of larger increases in environments other than the evolution environment. We also tested whether the increase in the evolution environment predicted the increase in other environments. A linear regression (not shown in figure) comparing the increase growth rate in the evolution environment to the mean increase in growth rate in other environments and another comparing the increase in maximum OD in the evolution environment to the mean increase in other environments showed no correlation for growth rate (spearman; r = -0.12, *p =* 0.19) but a significantly positive correlation for maximum OD (spearman; r = 0.35, *p =* 0.00015).

Finally, we ask whether the performance of a population in X environment following evolution in Y environment can be predicted by the performance of another population in Y environment following evolution in X environment. When we consider the composite fitness metric r-max X maximum OD, we found a significantly negative spearman correlation between the performance of Y evolved population in X environment and X evolved population in Y environment (Figure 5). This means that if a population evolved in environment X has large increases in environment Y it is more likely that a population evolved in environment Y will have smaller increases when assayed in environment X. It is probably not a coincidence that one of the major outliers to this trend (indicated by the red arrow in the figure 5) is the CA and CA no glucose environment combination. These two environments are probably highly correlated in the selection imposed meaning evolution in either environment is likely to result in increases in both environments, which is what we observe.

## Discussion

We found that fitness increases are dependent on the environment in which fitness is measured in addition to the environment in which evolution occurs. In some cases, we found that evolution can fail to increase fitness in the evolution environment while increasing fitness in alternative environments. In other cases, we see that the largest increases in fitness in some environments occur following evolution in other environments. Evolution could result in larger fitness increases in alternative environments for several reasons. It is possible that random non-selected mutations increase fitness in one environment and not the other, however this is probably unlikely as the chance of a random mutation having a large positive effect in any environment is low. A more likely explanation is that mutations that are advantageous in the evolution environment have larger beneficial effects in an alternate environment, or deleterious mutations that are able to accumulate in the evolution environment have a less deleterious effect in another environment or both.

Even if fitness isn’t changing, populations are still evolving due to new mutations and changes in allele frequencies (Tenaillon et al. 2016; Good et al. 2017). The rate at which deleterious mutations are introduced into a population may be countered by the rate at which beneficial mutations are introduced so that fitness does not change (Goyal et al. 2011). The beneficial mutations selected as evolution occurs in one environment may have a greater fitness effect in the other environment (Martin and Lenormand 2006), particularly if a population finds itself nearing the top of a fitness peak in the evolution environment. In fact, our results may in part be due to the decelerating rate of returns pattern that is often seen in evolutionary trajectories and is attributed to diminishing returns epistasis (Wünsche et al. 2017). If the ancestral genotype was already near or closer to the top of a fitness peak in some environments but not others it would likely constrain the changes in fitness in those environments (Schoustra et al. 2016; Wünsche et al. 2017) and result in the significant effect of assay environment that we observe. We can see this pattern in figure 3 indicating that fitness changes in an environment are at least in part dependent on the ancestral fitness in that environment.

**Figure 3.**
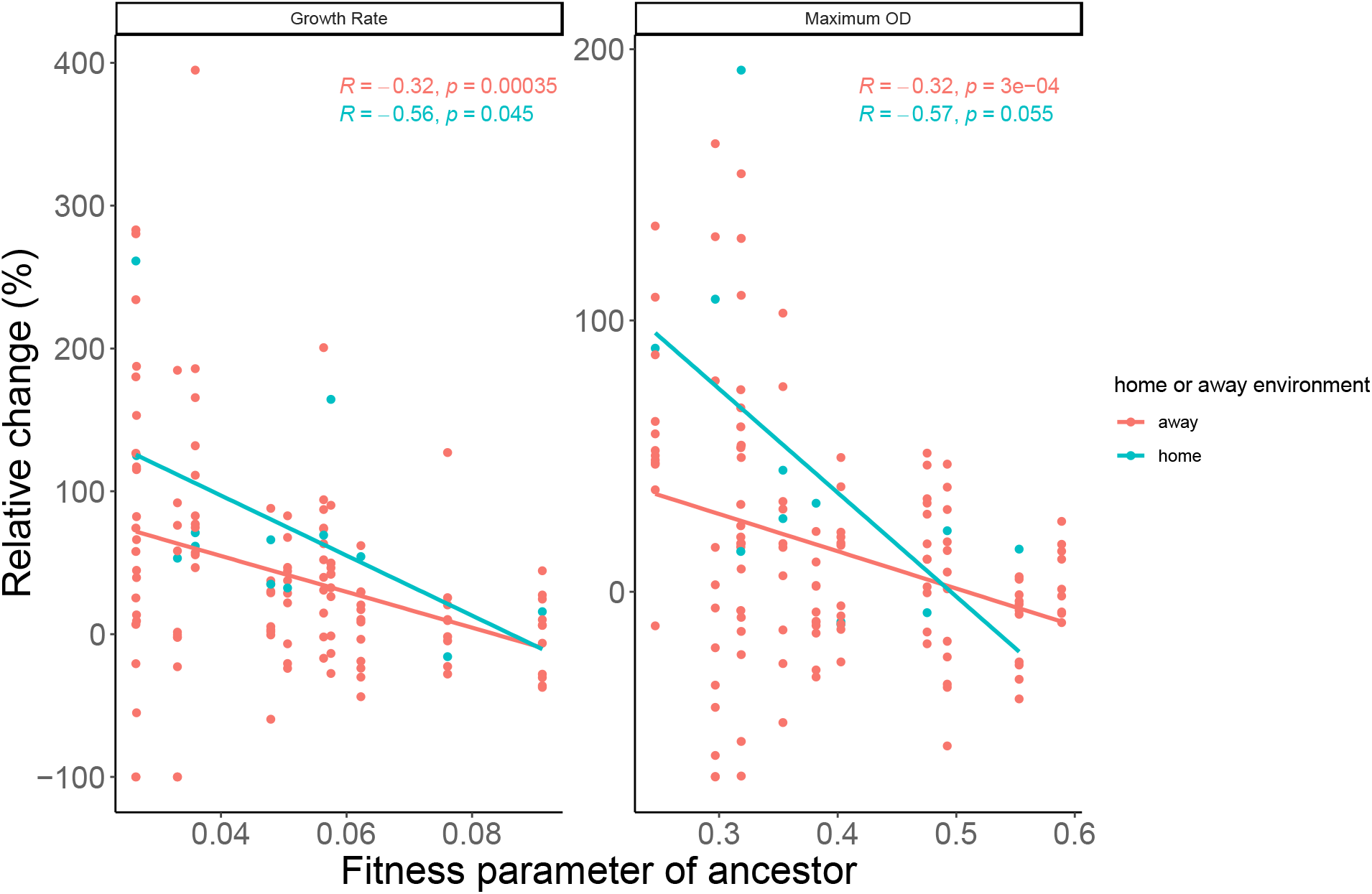
Relationship between ancestral fitness parameter and the change in that parameter following evolution. Each point shows an ancestral fitness parameter in one of 11 environments (x-axis) and the percent change in the fitness parameter in that environment (y-axis) following evolution in that environment (home; green points) or evolution in one of the 10 other environments (away; red points show mean of response following evolution in away environments). The growth rate parameter is shown on the left and the maximum OD650 is shown on the right. For each fitness parameter a linear regression is shown for the correlated and direct responses to evolution. In both cases lower ancestral fitness in an environment tends to results in larger fitness increases in that environment regardless of whether evolution occurs in that environment, however the relationship between ancestral maximum OD and the relative change in maximum OD was not significant when we considered only the direct response to selection.

On the other hand, a systematic effect of the evolution environment could result from mutations that fix in some environments also being beneficial in others (generalist mutations) while in other environments evolution fixes only specific mutations that are not beneficial in others (specialist mutations). However, this cannot explain how fitness in A could ever increase more after evolution in B compared to after evolution in A as we observe across temperatures or, for example, when fitness in acetate increases more after evolution in glycerol compared to after evolution in acetate. Fitness in A could increase more after evolution in B compared to after evolution in A if (i) some evolution environments experience different mutation rates so that evolution happens more quickly in some environments than in others, (ii) some environments select mutations that epistatically open up genotype space that allows higher fitness in alternate environments that is inaccessible when evolving in those environments, or (iii) there are many mutations with very small beneficial effect that are effectively neutral in one environment but of large enough effect to be selected in another so that more beneficial mutations in environment A accumulate when evolution happens in environment B.

It is important to remember that even very minor changes in the environment could result in these types of dynamics. For example, differences in the daily growth cycle and density range experienced could have profound effects (Li et al. 2018) so that whenever comparing the evolvability or the repeatability of evolution of different transfer regimes fitness should be measured under both regimes for populations from either regime.

We also found that the improvement of a population in an environment can be partially predicted by the improvement of a population in the reciprocal evolution and assay environment. If a population evolved in X has large increases in environment Y then a population evolved in Y is more likely to have small increases in environment X. Why this would be the case is not entirely clear. We might have expected this correlation (Figure 5) to be positive instead of negative if the selective pressure across the environments were highly correlated. The negative correlation that we observe could result if the selective pressures across environments are asymmetric, whereby evolution in X improves performance in Y but not vice versa.

**Figure 4.**
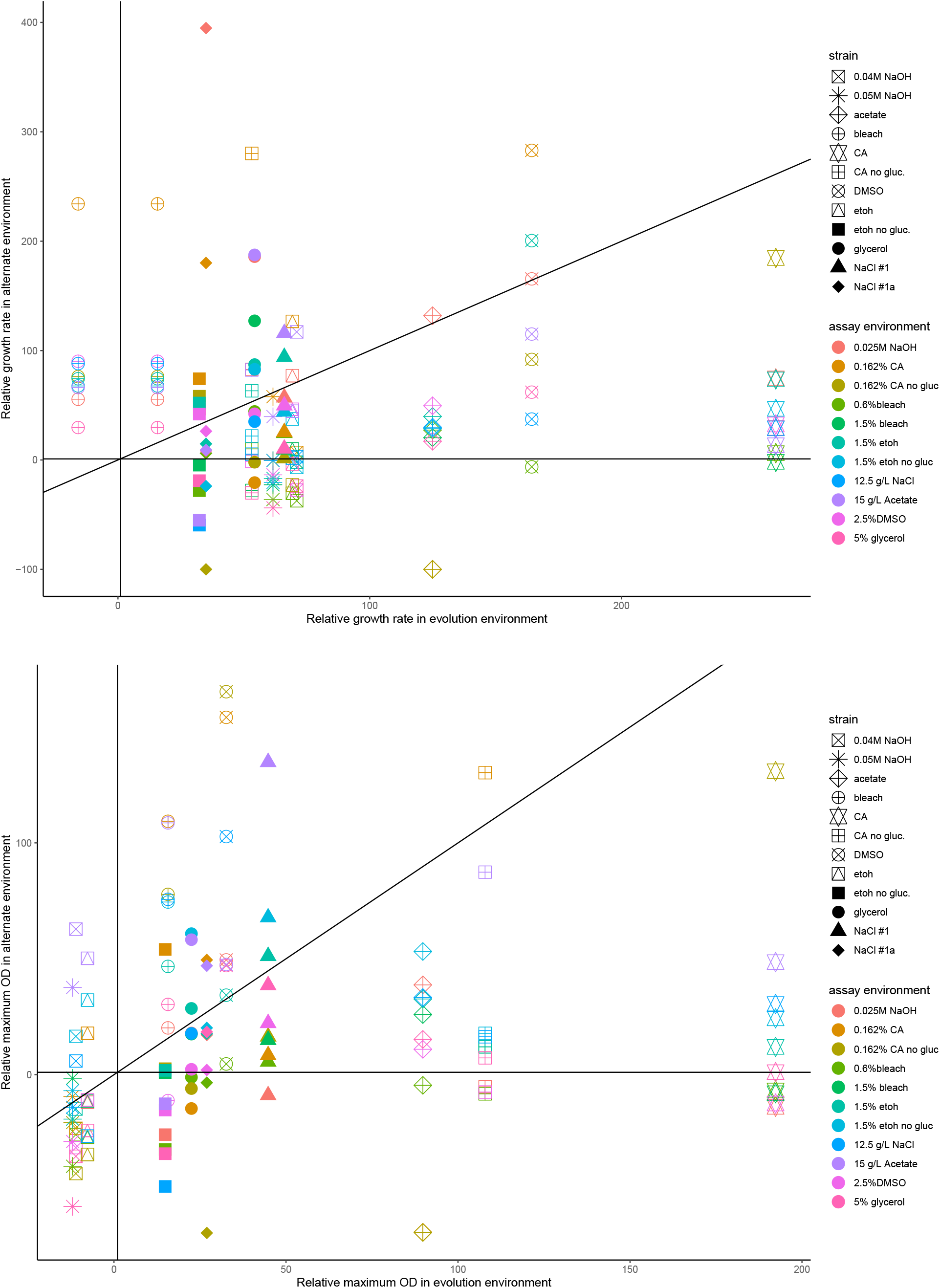
Trade-offs in relative fitness parameter in evolved vs. alternate environment. Each point shows the relative fitness parameter of an evolved population in its evolution environment (x-axis) and in one of the alternate environments (y-axis). The shape of the point indicates the population and its evolution environment while the color represents the alternate assay environment. A trade-off exists when points fall below 1 on the y-axis. If points are above the x=y line it indicates larger fitness increases in the alternate environment than the evolved environment. The top figure shows the mean growth rate while the bottom figure shows the mean maximum OD. The two evolution environments for the bleach evolved strain correspond to the two concentrations of bleach that were assayed (0.6% and 1.5%).

**Figure 5.**
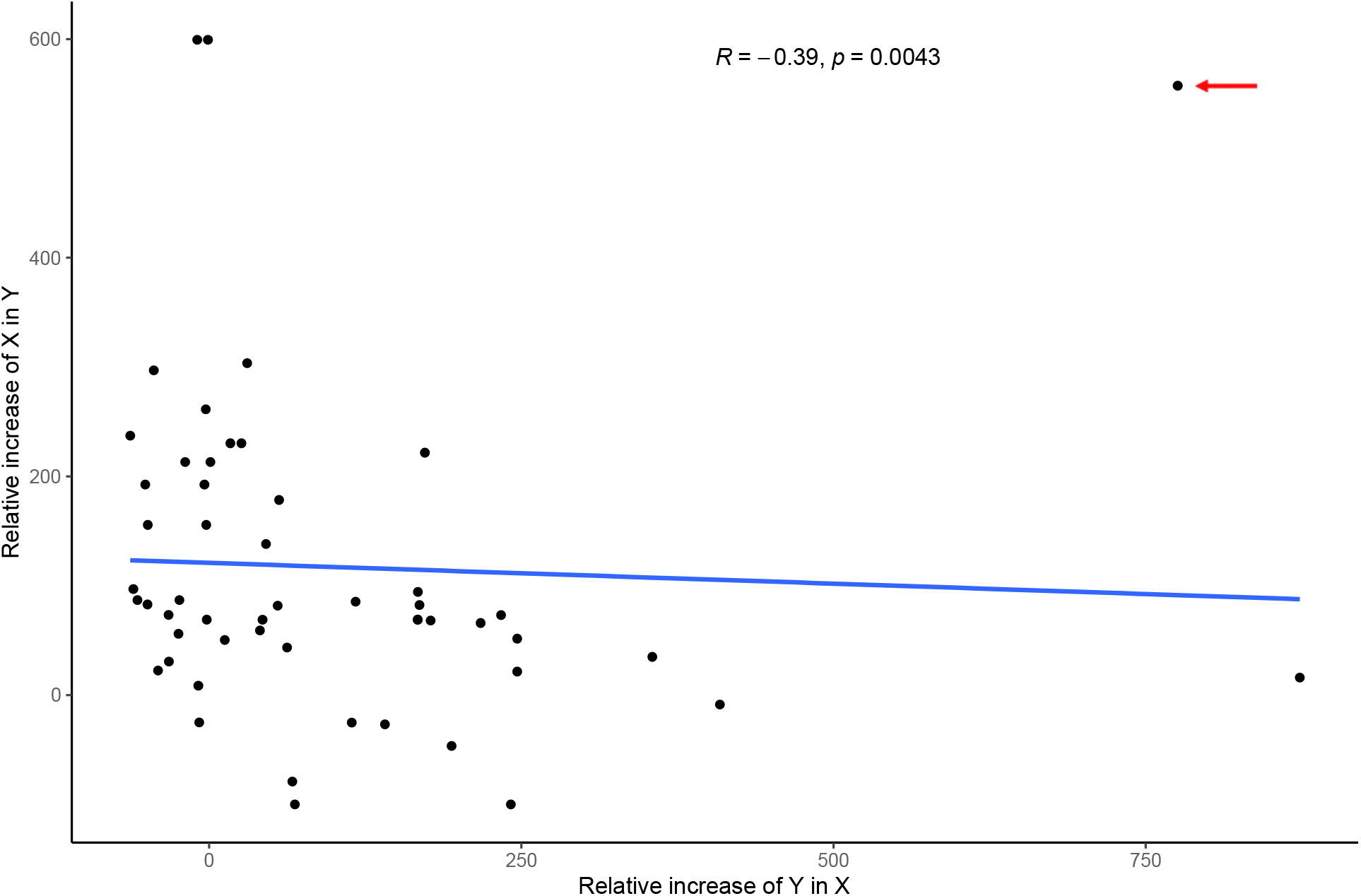
Asymmetry of the correlated responses. Each point shows one of 53 unique combinations of two environments. The y-axis shows the performance of a strain evolved in X and assayed in Y and the x-axis shows the reciprocal; the performance of a strain evolved in Y and assayed in X. Here performance is the relative increase in the composite fitness metric, r-max X maximum OD. Despite the outlying CA-CA no glucose point (indicated by the red arrow) we still see a significant negative spearman correlation.

## Conclusion

We show that the evolvability of a genotype in an environment is a combination of the adaptation that occurs in that environment and the way in which the environment constrains fitness. While these components are often treated as a single entity, they can also become decoupled if the correlated responses to evolution are measured. This can result in the interesting situation, in which evolution in environment A increases fitness in A more than evolution in B increases fitness in B while at the same time evolution in B results in greater fitness in A and B. In this case a genotype in environment A may appear more evolvable if we do not consider the correlated responses to evolution. However, when we consider the correlated responses, we can see that evolution in environment B promotes greater fitness increases in both environments and that fitness is simply more constrained in environment B. We show this situation across temperatures for the relative growth rate of *Tetrahymena* where the colder temperature behaves like environment A, although we note that this pattern is not consistent across all timepoints. We also show that this scenario is likely common across a variety of environmental pairs and that the fitness increases under a given environmental condition can be predicted by the ancestral fitness in that condition regardless of the environment in which evolution occurs. This result suggests there is a high degree of genetic correlation in fitness across environments but that fitness changes scale differently between environments.

